# Integrative network analysis interweaves the missing links in cardiomyopathy diseasome

**DOI:** 10.1101/2021.02.26.433014

**Authors:** Pankaj Kumar Chauhan, Ramanathan Sowdhamini

## Abstract

Cardiomyopathies are progressive disease conditions that give rise to an abnormal heart phenotype and are a leading cause of heart failures in the general population. These are complex diseases that show co-morbidity with other diseases. The molecular interaction network in the localised disease neighbourhood is an important step toward deciphering molecular mechanisms underlying these complex conditions. In this pursuit, we employed network medicine techniques to systematically investigate cardiomyopathy’s genetic interplay with other diseases and uncover the molecular players underlying these associations. We predicted a set of candidate genes in cardiomyopathy by exploring the DIAMOnD algorithm on the human interactome. We next revealed how these candidate genes form association across different diseases and highlighted the predominant association with brain, cancer and metabolic diseases. Through integrative systems analysis of molecular pathways, heart-specific mouse knockout data and disease tissue-specific transcriptomic data, we screened and ascertained prominent candidates that show abnormal heart phenotype, including *NOS3, MMP2* and *SIRT1*. Our computational analysis broadens the understanding of the genetic associations of cardiomyopathies with other diseases and holds great potential in cardiomyopathy research.

## Introduction

Cardiomyopathies are a severe and chronic health issue across the world^1–4^. These are complex heart muscle diseases compounded by genetic and environmental factors. Cardiomyopathies share overlapping genetic and phenotypic features with other diseases ^5–7^. Evidence from a growing number of studies suggests that several drugs, including anticancer, antiretroviral and antipsychotic, pose a potential risk of cardiotoxicity and drug-induced cardiomyopathies ^8,9^. It is well understood that sarcomere genes are the major drivers for cardiomyopathy phenotypes ^1,10,11^. Besides sarcomeric genes, cellular energy-related, Ca^2+^ handling and other genes are also implicated in cardiomyopathy ^10–13^. A subset of these genes influences cardiomyopathy expressivity and severity ^14,15^. We refer to such genes (variants) as ‘modifier genes’ that influence the function of another gene. Hamilton et al. defined modifier genes as the genetic variants that can change the phenotypic outcome of an independent ‘conditioning’ variant at another locus ^16^. Genome-wide association studies (GWAS) and candidate gene analysis approaches point to many common genetic variants as one of the plausible reasons for toxicity or induced cardiomyopathy ^17–19^. Hence, a fundamental knowledge of the cardiomyopathies and the molecular players involved is critical for developing novel approaches for its prevention and treatment. Modifier genes are suitable candidates for finding biomarkers ^20,21^, drug toxicity explanations ^22,23^, and possible missing links between distinct diseases.

The emergence of network biology has allowed us a more refined understanding of complex systems like protein-protein interactions and disease-disease links ^24,25^. Disease network analysis is helpful in disease epidemiology as it facilitates and projects a simple concept of the relative risks of diseases and characteristics of their shared architecture ^26–28^. In this direction, the first genotype-based human disease network was constructed based on commonly shared genes between diseases. It was a novel attempt to show the global organization of diseases and functional modules ^29^. Now, this methodology has been used in therapeutic innovation and disease drug repurposing ^30^. Cardiomyopathies have been reported in several disease network studies ^29,31^. However, to the best of our knowledge, none of the studies exclusively focus on cardiomyopathies, potential modifiers and their associations, consequently requiring in-depth, rigorous analysis. In this perspective, our study is the first of its kind large-scale exploration of candidate (potential modifier) genes and disease associations involved in cardiomyopathy.

The current study employs an integrative systems biology approach to understand cardiomyopathy-centric complexities and the shared genetic architecture. Firstly, we performed comprehensive mining of publicly available genetic, protein-protein interaction (PPI), mouse phenotype, and transcriptome data related to cardiomyopathies. Secondly, we generated a cardiomyopathy-centric diseasome network based on genetic data. Thirdly, we explored human interactome datasets to predict candidate (potential modifier) genes in cardiomyopathies and assessed their associations with other diseases. Furthermore, we screened our findings using mouse-abnormal heart phenotype data and transcriptome datasets from the European Nucleotide Archive (https://www.ebi.ac.uk/ena/browser/home) repository to associate the cardiomyopathy-centric candiate genes to other disease phenotypes.

## Results

### Genetic connectivity of cardiomyopathy genes with other pathophysiological diseases

To identify the molecular players underlying genetic links between cardiomyopathies and other diseases, we first constructed a comprehensive diseasome network based on the genetic data from publicly available datasets (see Methods). We extracted a non-redundant set of 4,406 disease phenotypes and associated genes in this process. Furthermore, each disease was partitioned into disease-category after merging similar diseases using fuzzy matching. ultimately reducing the size to 2722 distinct disease-gene associations (*see Table S1A*). Next, to construct cardiomyopathy-centric diseasome, we investigated only those disease-gene associations that consisted of at least one cardiomyopathy gene(*see Table S1B*). This exercise resulted in a bipartite network with 146 diseases and 1,929 genes (*see Table S1C*). Ultimately, this bipartite network was projected to the disease network based on common genes. This disease-projected network was termed as cardiomyopathy-centric diseasome consisting of 146 diseases with 1193 distinct links (*Figure 1*).

**Figure 1.**
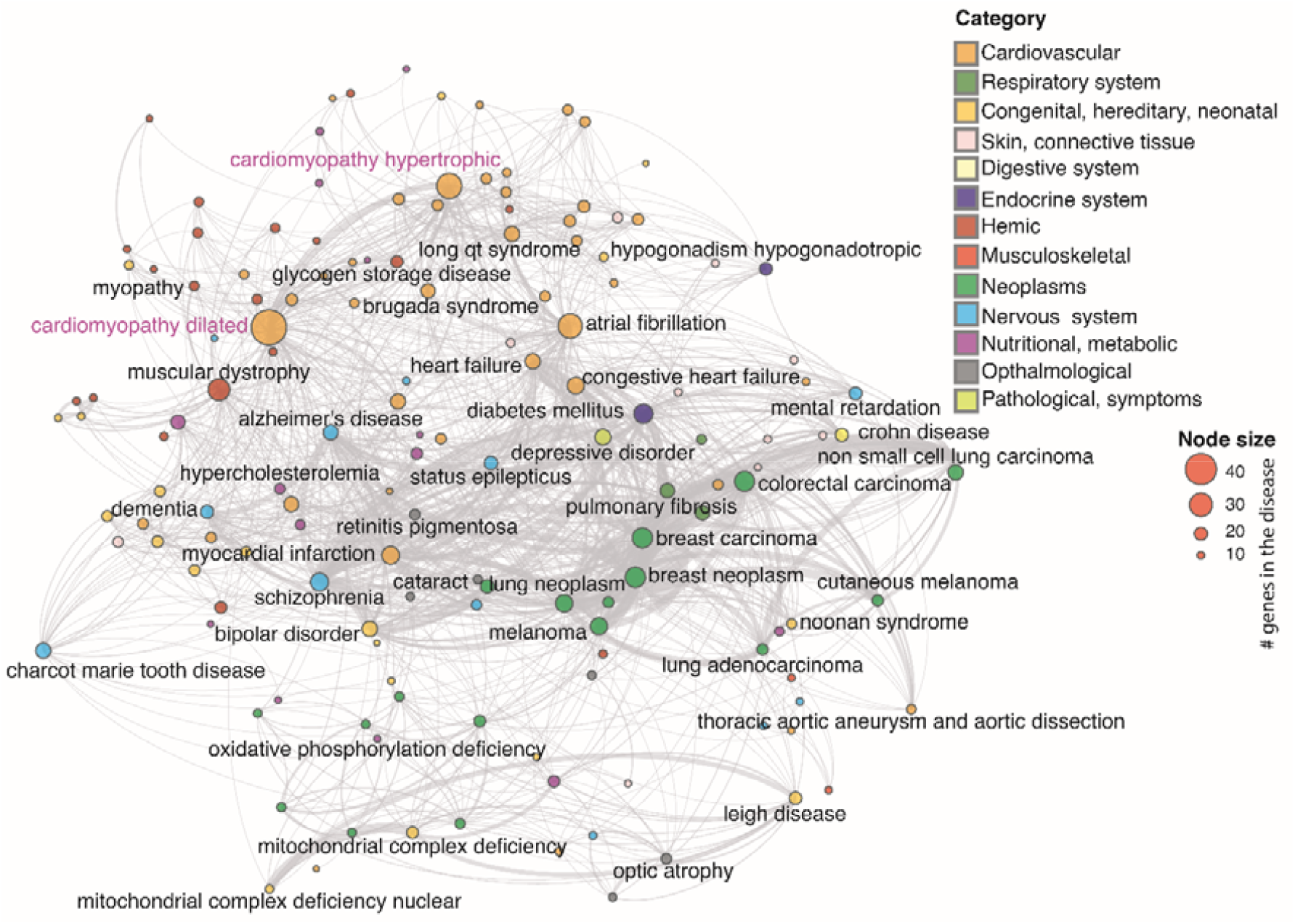
Cardiomyopathy-centric diseasome: A graph representing the cardiomyopathy-centric diseasome network, in which two diseases are linked if a gene is implicated in both diseases. It is constructed by projecting a cardiomyopathy-centric bipartite graph. Each node represents a distinct disease in the network, and it is coloured based on the disease category it belongs to. The size of each node is proportional to the number of genes in that particular disease. The name of diseases with > 20 associated genes are labelled in the network. There are 12 disease categories in the cardiomyopathy-centric diseasome as labelled in the legend. The links between disease pairs are shown in grey colour. The weight of a link is proportional to the number of genes implicated in both diseases.

### Evaluation of the cardiomyopathy-centric diseasome network properties

Cardiomyopathy-centric diseasome network revealed that cardiomyopathies were linked with cardiovascular, neoplasms, musculoskeletal, metabolic and nervous system disease-categories (*Figure 1*,*2A*). The predominant category was the cardiovascular system occupying 28.7% of the total associations, followed by musculoskeletal and congenital, each sharing 13.7% interactions. Surprisingly, neoplasms were also linked to cardiomyopathies (12.2%) dominated by the RAF1 gene (41 %). Metabolic disorders were another significant contributor (10.0%). We performed network statistics such as degree, betweenness and closeness centrality, degree distribution (k) and gene distribution on the cardiomyopathy-centric diseasome network (*see Table S2*). The degree distribution points that most diseases are linked to only a few other diseases (*Figure 2B-C, also see Figure S1*). In contrast, intended cardiovascular diseases such as DCM (k=96) and hypertrophic cardiomyopathy (HCM) (k=63) are linked to many diseases. For comparison of cardiomyopathic-centric diseasome with random control, we reshuffled the genes of each disease (10,000 trials). Results show that cardiomyopathic-centric diseasome has significantly higher disease links (z-score = 6.652, p-value = 1.44e-11) than random expectation. Also, most of the intra-disease category links are significantly high compared to the one in the random networks (*Figure 2D-E)*.

**Figure 2.**
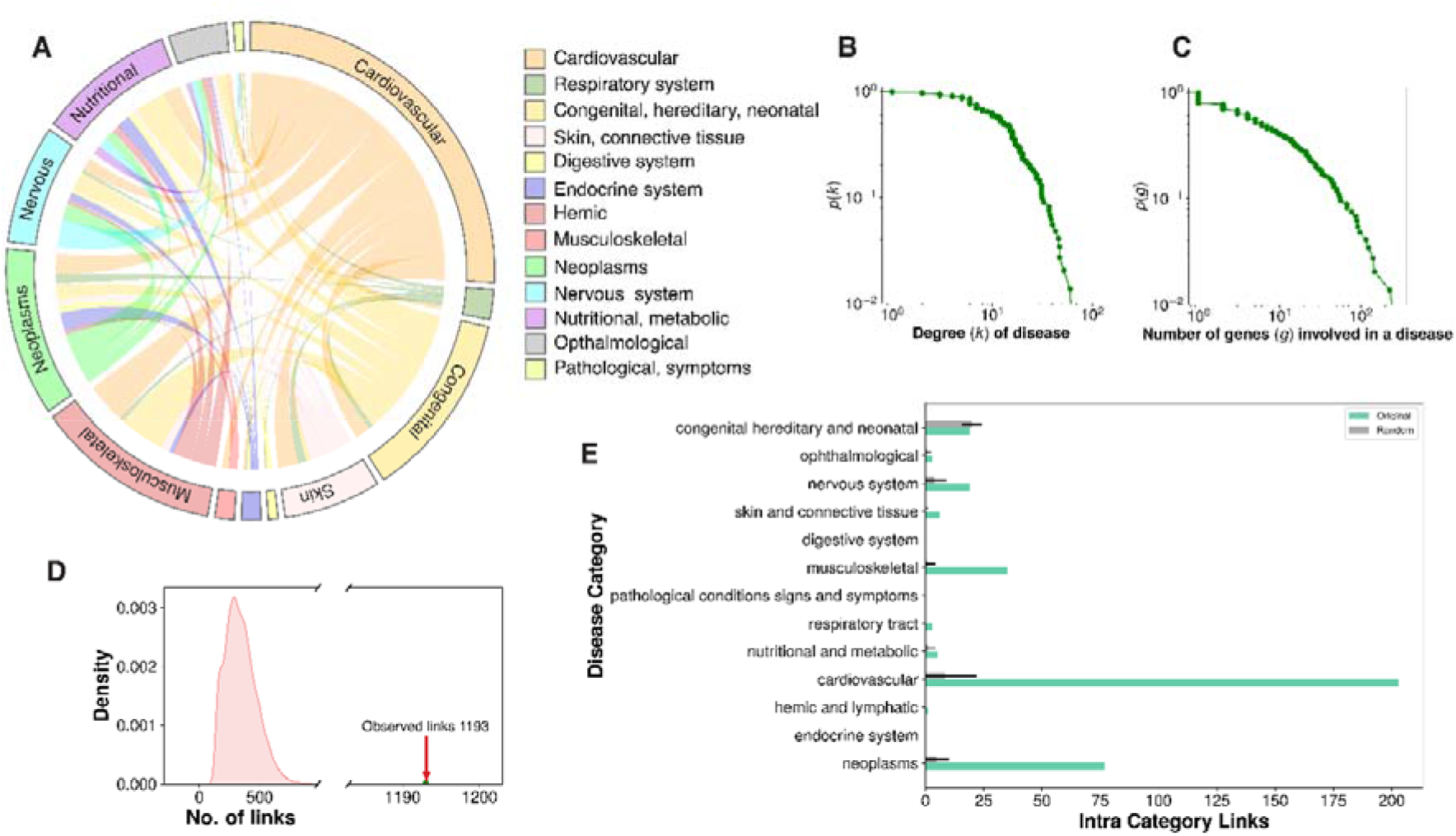
Diseasome properties: Overview of the cardiomyopathy-centric diseasome network. A) Circos plot showing the inter disease category association of diseases in the cardiomyopathy-centric diseasome network. The names of the 12 disease classes are shown on the right and coloured accordingly. B) Distribution of the degree k (number of diseases sharing common genes) in cardiomyopathy diseasome. The green dots represent the logarithmically binned data. C) Distribution of the gene size g (number of genes implicated in a particular disease) in the cardiomyopathy-centric diseasome. D) Number of disease links in the cardiomyopathy-centric diseasome versus random expectation. E) Intra-disease category links distributions for real data and random trials. Green bars represent real data while gray bars show average intra-disease category links in the random trials. Error bars denote standard deviation.

Functional diversity of cardiomyopathy genes can be critical in the risk of cardiotoxicity and phenotypic modulation of cardiomyopathies. We conducted a suite of analyses to statistically quantify biological and functional diversity in the cardiomyopathycentric diseasome network using the molecular pathways homogeneity (PH) and gene ontology homogeneity (GH) ^29^ distributions (*Figure 3*). We also performed randomization of genes in each pathway and GO terms to evaluate the significance of real distribution v/s random simulations using normal distribution test statistics (z-score and P-values, see Methods). We observed that disease-associated PH are significantly higher at perfect homogeneity value in comparison to the random control (fold change = 19.7, z-score = 23.31, p-value =1.75e-120). Similar trend was observed in gene ontology homogeneity (GH) analysis in all three branches (molecular function fold change = 7.9, z-score = 9.3, p-value =7.02e-21; cellular components fold change = 2.5, z-score = 5.6, p-value =1.07e-08; and biological processes fold change = 12.5, z-score = 13.7, p-value =5.07e-43; respectively). Disease degree vs PH or GH distribution showed that higher PH or GH values showed decline in the disease degree. Likewise, distribution of common shared genes between two diseases and their average PH followed the similar trend. These results point that diseases with higher number similar functional genes tend to have few disease association. On the whole, pathway and gene ontology homogeneity analysis demonstrated that disease genes tend to be connected functionally and this property can be explored to predict new genes.

**Figure 3.**
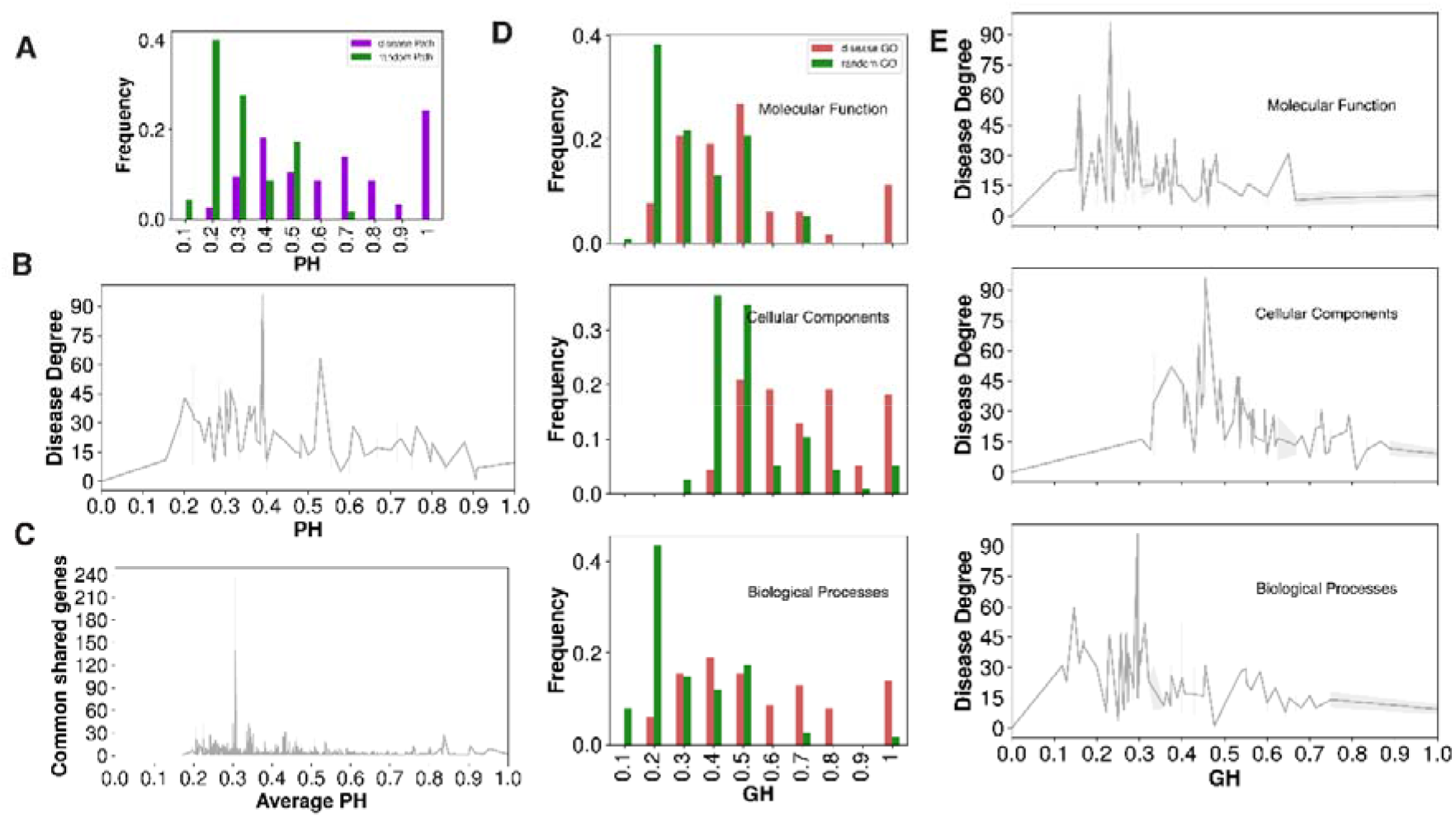
Diseasome properties: A) A distribution of the molecular pathways homogeneity (PH) of individual diseases in cardiomyopathy-centric diseasome. Purple bars represent the actual diseasome, and the green bars show the random control calculated for each disease by randomly choosing the same number of genes. The PH is formulated as the maximum possible fraction of genes associated with the same molecular pathway in individual disease. B) A lineplot of disease degree versus PH value distribution in the diseasome. C) The distribution of number of common genes between two disease phenotype and their average PH value in the cardiomyopathy-centric diseasome. D) The GO homogeneity measure of cardiomyopathy-centric diseasome genes for all GO categories -cellular component (CC)(Middle), biological process (BP) (Bottom), and molecular function (MF) (Top). The GH for each disease term was calculated like the PH analysis. Red bars represent the actual diseasome histogram, and the green bars show the random control. E) A lineplot of disease degree and GH value distribution for each GO category in the diseasome.

### Identification of candidate genes in cardiomyopathies and disease modules

Further extending the clustering of genes visible in the PH and GH analysis of cardiomyopathy-centric diseasome, we predicted new candidate (potential modifier) genes in cardiomyopathies. For this, we utilized the DIseAse MOdule Detection (DIAMOnD) algorithm on network topology created by the human interactome (see Methods) ^32^. The DIAMOnD algorithm is a popular method used in several studies for disease-gene prediction ^33–35^. It explores the topological neighbourhood of the seed genes in the human interactome and identifies newer genes based on significant connectivity to the seed genes (see Methods) ^32^. An exhaustive number of DIAMOnD genes (1000 candidate genes) were predicted for each cardiomyopathy. Since the DIAMOnD algorithm continues to identify and associate newer genes to the initial set of seed genes, it is necessary to set a limit for the expansion of such association in the entire gene dataset. For this, we quantified the biological relevance of the newly predicted gene using the molecular pathway data (see Methods). We tabulated all the molecular pathways enriched in pathway enrichment of seed genes (adjusted p-value = 0.05) for individual cardiomyopathies. Subsequently, DIAMOnD genes showing enrichment with same pathways were considered true hits for candidate genes. We profiled approximately the first 601, 508, and 31 DIAMOnD genes that show a clear and significant biological association with HCM, DCM, and arrhythmogenic right ventricular cardiomyopathy (ACM), respectively (*Figure 4A-F*). In minor forms like idiopathic cardiomyopathy (IdCM), hypertrophic obstructive cardiomyopathy (HoCM), restrictive cardiomyopathy (RCM) and HcCM, we found 20, 10, 7, and 1 DIAMOnD genes, respectively, while in amyloid cardiomyopathy (AmCM), mitochondrial cardiomyopathy (MtCM), diabetic cardiomyopathy (DbCM) boundary was not possible due to a limited number of seed genes. To further filter the candidate genes associated with cardiomyopathies, we verified if their ortholog genes in the mouse knockout dataset showed abnormal heart phenotype (see Methods). Only mapped candidates were used for further analysis. This result led to the identification of 53, 45 and 2 mapped candidate genes in HCM, DCM, and ACM, respectively (*Figure 4G-I*, and *also see Table S3*). IdCM, HoCM, RCM, and HcCM DIAMOnD genes did not show any overlapping genes with mouse phenotype data (*see Figure S2*). Further, to compare our constructed human interactome, we considered two independent datasets (HuRI and BioPlex3) ^36,37^. We looked for the interaction of human ribosomal proteins. Both of these datasets failed to account for interactions in the ribosomal complex. However, these interactions were present in our human interactome and the STRING database ^38^ (*see Figure S3*). We, therefore, continued with our interactome only.

**Figure 4.**
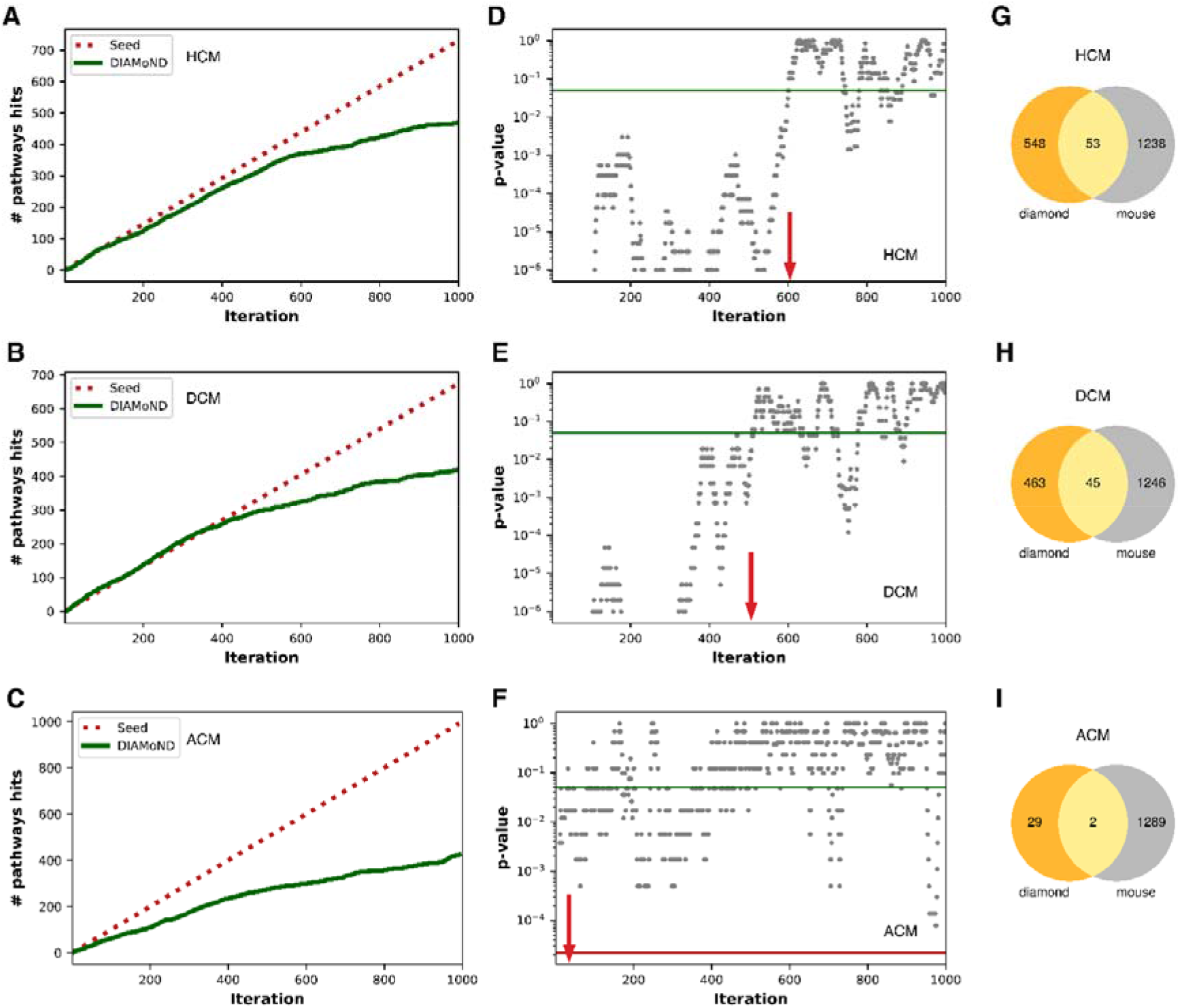
Candidate genes: Graph illustration of the biologically relevant association of the predicted DIAMOnD genes and filtering based on abnormal heart phenotype in the mouse. (A,B,C) Panels correspond to true molecular pathways hits of DIAMOnD genes of hypertrophic cardiomyopathy (A), dilated cardiomyopathy (B), and arrhythmogenic cardiomyopathy (C), respectively (red lines, seed genes; green lines, DIAMOnD genes). (D,E,F) Panels show the distribution of p-value for DIAMOnD genes where a sliding window of DIAMOnD genes size equal to seed genes was used for Fisher’s test for each iteration. The cut-off of p-value = 0.05 (green line) was chosen for selecting DIAMOnD genes in each cardiomyopathy. Red arrow marks the boundary of reliable DIAMOnD genes (G,H,I) Panels are Venn diagrams that project the overlap of DIAMOnD genes and mouse genes showing abnormal heart phenotype.

### Candidate genes fill the missing links in cardiomyopathy-centric diseasome

Next, we assessed the role of these candidates or potential modifiers in the expansion of the cardiomyopathy-centric diseasome. Apart from the original 269 cardiomyopathies associations to other diseases, the inclusion of predicted candidate genes to cardiomyopathies leads to an additional 221 links to other diseases (*see Table S4*). Interestingly, apart from heart-related diseases and cancers, diabetes, rheumatoid arthritis, lipodystrophy, nonalcoholic fatty liver disease, brain disorders (like Alzheimer’s, depressive disorder, schizophrenia, and mental retardation) were part of cardiomyopathy associations due to the common candidate genes.

The notable candidates associated with these diseases were *IL6, NOS3, MMP2, SIRT1, CAV1* and *ESR2* (*Table 1, also see Table S4A*). We observed that these candidates contributed to 22.62 %, 11.31%, 9.76%, 5.65%, 5.65% and 5.14% of the new disease association with cardiomyopathies, respectively *(see Table S4B)*. We examined the RNA expression of the modifier genes in the primary tissues to associate whether candidates are indeed expressed in the heart and other tissues. We explored the Human Protein Atlas (HPA) dataset for finding gene expression of candidates for this analysis ^39^. Except for *TBL1Y, PAX5* and *ESR2*, most candidates showed reliable and abundant expression in the HPA tissues. The RNA expression of important candidates like *IL6, NOS3, MMP2, SIRT1* and *CAV1* was observed in the heart muscle tissue *(see Table S4C)*.

**Table 1:** A representative disease-disease association of predicted candidate genes that are screened based on mouse phenotype genes

| S. No. | Disease A | Disease B | Candidate genes |
| --- | --- | --- | --- |
|  | Hypertrophic cardiomyopathy | Alzheimer's disease | *NOS3* |
| 1 | Alzheimer's disease | Dilated cardiomyopathy | *NOS3* |
| 2 | Dilated cardiomyopathy | Non-alcoholic fatty liver disease | *SIRT1* |
| 3 | Hypertrophic cardiomyopathy | Non-alcoholic fatty liver disease | *SIRT1* |
| 4 | Dilated cardiomyopathy | Rheumatoid arthritis | *IL6, MMP2, IL2RA* |
| 5 | Dilated cardiomyopathy | Diabetes mellitus | *NOS3, IL6, SIRT1, IL2RA* |
| 6 | Dilated cardiomyopathy | Lipodystrophy | *CAV1* |
| 7 | Dilated cardiomyopathy | Schizophrenia | *IL6, ESR2* |
| 8 | Hypertrophic cardiomyopathy | Depressive disorder | *IL6, ESR2* |
| 9 | Hypertrophic cardiomyopathy | Diabetes mellitus | *NOS3, IL6, SIRT1, IL2RA* |
| 10 | Hypertrophic cardiomyopathy | Schizophrenia | *IL6, ESR2* |
| 11 | Hypertrophic cardiomyopathy | Parkinson disease | *IL6* |
| 12 | Hypertrophic cardiomyopathy | rheumatoid arthritis | *IL6, MMP2, IL2RA* |
| 13 | Dilated cardiomyopathy | Parkinson disease | *IL6* |
| 14 | Hypertrophic cardiomyopathy | Lipodystrophy | *CAV1* |
| 15 | Dilated cardiomyopathy | Unipolar depression | *IL6* |
| 16 | Hypertrophic cardiomyopathy | Mental depression | *IL6* |
| 17 | Hypertrophic cardiomyopathy | Intellectual disability | *RAC1* |
| 18 | Dilated cardiomyopathy | Bipolar disorder | *IL6* |
| 19 | Dilated cardiomyopathy | Intellectual disability | *RAC1* |
| 20 | Hypertrophic cardiomyopathy | Bipolar disorder | *IL6* |

We extended our study of candidate genes identified in cardiomyopathy to establish a connection with non-cancer and non-heart related diseases. Our analysis predicts that IL6 is a potential gene relevant to the association of bipolar disorder, depressive disorder and schizophrenia with DCM. Similarly, *NOS3* connects Alzheimer’s disease and Diabetes mellitus with HCM and DCM. On the other hand, *MMP2* is implicated in the association of rheumatoid arthritis to HCM and DCM. *SIRT1* is found to connect diabetes mellitus and non-alcoholic liver disease to HCM and DCM. Lastly, *CAV1* is involved in relating lipodystrophy to HCM and DCM.

To further support identified candidate genes, we analysed disease tissue-specific RNA-seq datasets from the ENA repository (see Methods). However, we could not find disease tissue-specific data for many diseases. So we considered only diseases whose data was available as the case studies. *IL6, NOS3, MMP2* and *SIRT1* were implicated in these diseases. The results showed that most candidate genes are expressed in the tissues of interest. We could not find sufficient expression of *IL6* in our datasets, so we ignored it. Our results revealed that *NOS3* is expressed in both Alzheimer’s and DCM. Similarly, *SIRT1* was expressed in HCM, non-alcoholic fatty liver disease and Diabetes mellitus. Further, *MMP2* expression was observed in HCM and rheumatoid arthritis (*see Figure 5B-D*).

**Figure 5.**
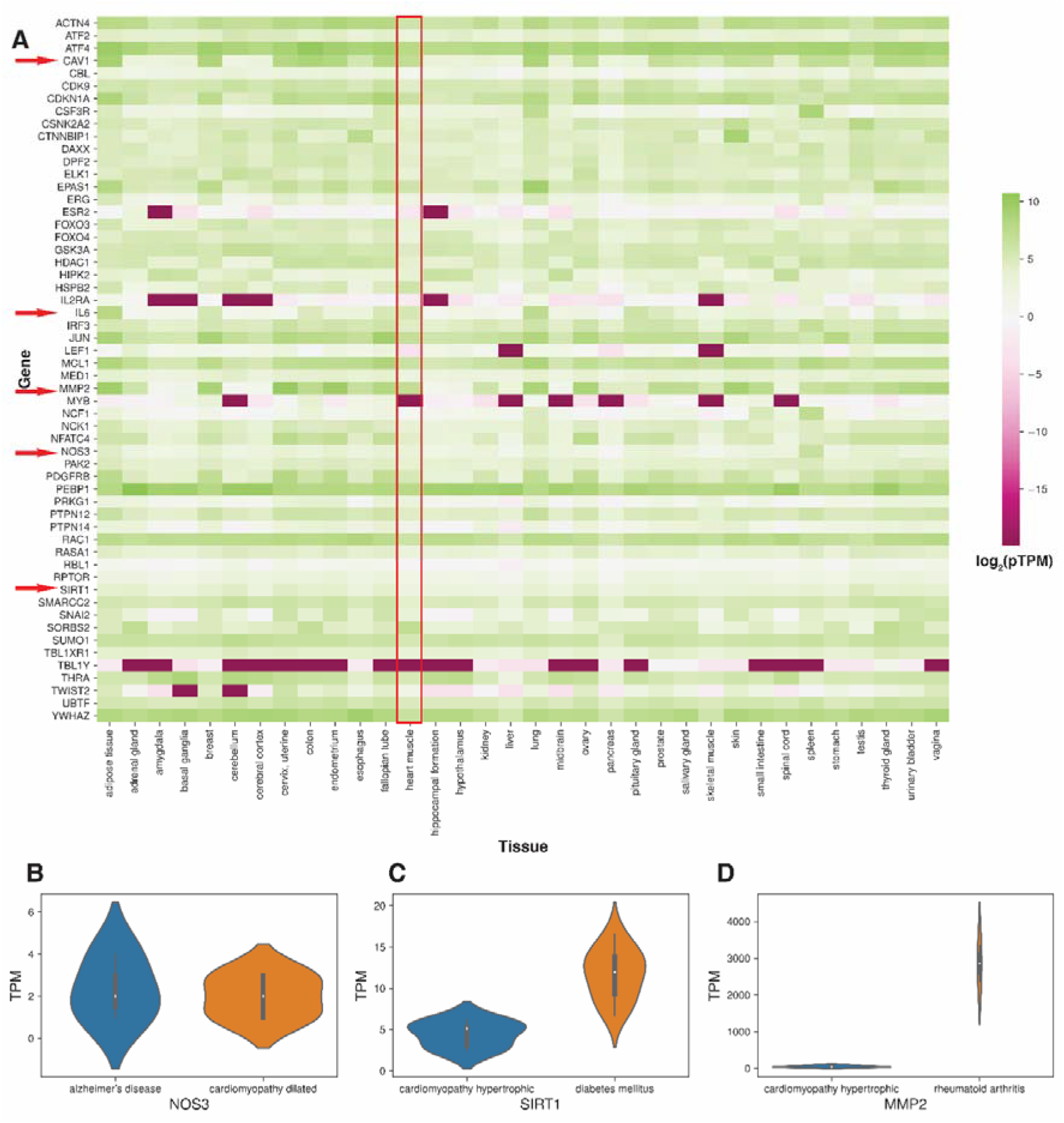
Tissue expression of candidate candidates: Overview of the tissue specific RNA expression of the candidate genes in cardiomyopathy-centric diseasome. A) The protein-transcript per million (pTPM) expression values of candidate genes across major tissue into GTEx dataset of the Human Protein Atlas database. In the heatmap, values are log2 transformed with green color showing abundant transcripts while pink color depicting extremely low expressed transcripts. Red arrows point to important candidates observed in the study. Heart muscle tissue is highlighted with red box. B) A violinplot shows the expression of NOS3 gene in Alzheimer’s and dilated cardiomyopathy. Transcript per million (TPM) of NOS3 derived from Brain and heart tissues (PRJEB28518 and PRJNA494688) were used. C) Tissue specific expression of SIRT1 gene in hypertrophic cardiomyopathy and Diabetes mellitus. Expression of SIRT1 (TPM) was derived from dermal endothelial and heart tissues (PRJNA352990 and PRJNA358470). D) MMP2 gene expression in rheumatoid arthritis and hypertrophic cardiomyopathy from SRP009315 and PRJNA352990 datasets.

NOS3 (Nitric Oxide Synthase 3) produces the gasotransmitter nitric oxide (NO) that is implicated in vascular smooth muscle relaxation through a cGMP-mediated signal transduction pathway ^40^. Polymorphisms in the *NOS3* gene have been implicated in dilated cardiomyopathy ^41^. SIRT1 (Sirtuin 1), a crucial multi-functional protein mainly expressed in the endocrine tissues, is involved in controlling DNA repair, tissue regeneration, cell survival, inflammation, signalling, and circadian ^42^. Its deficiency leads to progressive dilated cardiomyopathy in adult mice ^43^. Also, PPARα-Sirt1 complex attributes to cardiac hypertrophy and failure by suppressing the ERR transcriptional pathway ^44^. MMP2 (Matrix Metalloproteinase 2) protein is known to degrade collagen, elastin, fibronectin, gelatin, and laminin and has both proinflammatory and anti-inflammatory roles in numerous tissues ^45^. Its expression is altered in skeletal muscle during heart failure and diabetic cardiomyopathy in rates ^46,47^.

We observed that *NOS3, SIRT1* and *MMP2* are prominent candidate genes. Further, these genes can be subjected to experimental validation. Similarly, this approach paves the way for candidate genes in a larger tissue-specific disease association studies.

## Discussion

Cardiomyopathies are important age-related diseases. Genetic, environmental and dietary factors contribute to disease severity; however, much remains to be discovered. Here, we applied an integrative systems biology approach to analyse human genetic, molecular interactome and transcriptome data to elucidate the molecular players involved in cardiomyopathy-centric diseasome. In cardiomyopathy research, no such attempts have been made previously involving genetic connectivity, protein interactions and tissue expression. Our cardiomyopathy-centric diseasome connected to many pathophysiological diseases from neurological, musculoskeletal, metabolic and neoplasms. These associations are constructed using datasets of already reported genetic studies and hence could play a significant role in deciphering drug-toxicity, off-target binding and/or novel targets in such diseases.

Our analysis revealed that the molecular players in a specific disease are generally clustered (i.e. form disease modules) supported by means of PH and GH assessment. Such genes carry out similar functions or are part of similar cellular processes. Notably, based on PH and GH assessment, disease genes were far more homogenised than the random control, suggesting a bona fide clustering of such genes involved in a disease. We expanded cardiomyopathy involved (seed) genes using DIAMOnD and assessed for biological relevance using the molecular pathways dataset (MSIgDB). Additionally, the mouse phenotype dataset helped to screen candidate (potential modifier) genes using IPMC phenotype data for heart-related abnormalities. Our analysis identified a set of candidate genes across different cardiomyopathies (HCM, DCM and ACM). We looked at the prominent diseases associated with these candidates. Our structured enquiries highlighted the notable candidates involved in linking cardiomyopathies to non-alcoholic fatty liver disease, diabetes mellitus, rheumatoid arthritis, as well as brain disorders like Alzheimer’s, bipolar disorder, mental retardation, schizophrenia, and depressive disorders.

Tissue-specific transcriptome datasets could be of vital use in discovering candidate genes. We used HPA and ENA RNA-seq) data to associate candidate genes from cardiomyopathy to other diseases. Interestingly, we found that the majority of the genes were abundant in major tissues, including heart muscle. We showed that *NOS3, CAV1* and *SIRT1* are expressed in the cardiomyopathies as well as other non-cancer and non-heart related diseases. Such clues arising from diseasome network analysis can benefit in drug repurposing studies and assist in finding possible off-targets and hidden genetic overlaps between cardiomyopathies and other co-morbidities.

In summary, unlike previous studies that provide a limited description of cardiomyopathies from the purview of the diseasome network, the current study focuses on the key molecular entities involved in these heart diseases. Our integrative network-driven approach predicts and highlights the prominent candidate (potential modifier) genes that associate cardiomyopathies with Alzheimer’s disease, rheumatoid arthritis, diabetes and non-alcoholic fatty liver disease. Our approach demonstrates the power of network study to find such novel candidate genes. Furthermore, this study points to the molecular candidates that can be a target for developing effective therapies against cardiomyopathies, apart from providing us with the molecular markers for such genetic disorders. Additionally, similar work can be performed for other disease-centric networks as well.

### Limitations of the study

This study strongly relies on disease genes association data as well as human interactome data. The primary limitation of this study is the incompleteness of initial data. Further, each dataset may contain noise, which may remain in the outcome even after processing. Methods employed for this very challenging problem of automated full-text analysis in large-scale data are limited in accuracy. Further, this study is predictive in nature, and experimental validation (such as cell assay, RNAseq) is beyond the scope of the current study. Nevertheless, cardiomyopathy-centric diseasome and candidate genes prediction would facilitate a comprehensive understanding of cardiomyopathies.

## Methods

### Statement on Data

Authors from this study reporting experiments on human data, human genome data and/or the use of human tissue samples confirm that all experiments were performed in accordance with the relevant guidelines and regulations.

#### 1 Data sources and processing

Publicly available gene and disease association data were downloaded from OMIM(v2018), ClinVar(2020), HumSaVar(2020) and DisGeNet(2020) datasets (Amberger et al. 2015; Landrum et al. 2018; Piñero et al. 2020). From the OMIM dataset, only ‘(3)’ marked diseases were selected for curated data. Similarly, in DisGeNet, diseases with a score above 0.4 were considered. HUGO Gene Nomenclature Committee (HGNC) gene symbols were assigned to each gene for a consistent and accurate name ^51^. Synonymic or alternative gene names were reduced to the HGNC gene primary symbol, as reported in HGNC (June 2020 release). All disease-gene datasets were merged and manually curated for a comprehensive non-redundant disease-gene data. Similar diseases, disease sub-types or the same disease with different names were merged (e.g. d-2 hydroxyglutaric aciduria and l-2 hydroxyglutaric aciduria were merged and named hydroxyglutaric aciduria) using fuzzy-wuzzy module in python. MeSH 2020 (2020), GARD (2020), and Literature data were integrated to categorise diseases. The category names were assigned according to significant organ systems based on MeSH 2020, and similar types were merged, thus restricting the number of the category to 12. A different ‘multiple’ group was created to cater to diseases belonging to multiple categories. The dominant type was assigned if a disorder was part of many categories. In the case of multiple dominant categories, it was assigned to the ‘multiple’ category. For human interactome construction, PPI data (HPRD, MINT, and IntAct) were integrated with other interactions (e.g. protein complex and kinase substrate) from CORUM and Phosphositeplus as well as transcription factors from the TRRUST database ^52–57^. For checking biasness analysis of interactome HuRI, BioPlex3 and STRING datasets were used ^36–38^. Custom python and R scripts were used for the analysis (available at http://caps.ncbs.res.in/download/cardiomyo_diseasome).

#### 2 Diseasome construction and network analysis

A bipartite network was constructed from the previous disease-gene data with diseases as one category of nodes and genes as another type. A disease-gene pair was linked if the gene was part of that disease. The diseasome was constructed by projecting the bipartite network on disease nodes. Further cardiomyopathy-centric diseasome network was obtained by retaining only those diseases in which at least one cardiomyopathy gene was common. Network analyses like degree distribution of diseases and distribution of genes in disease were carried out in the networkx module of python. Random networks (diseasome) were generated by shuffling the genes in the original diseasome. For each disease, an equivalent number of randomly shuffled genes were assigned and diseasome was constructed based on the common shared genes between two diseases. Statistical significance of cardiomyopathy-centric diseasome was calculated from 10000 trial of random networks. Normal distribution was used for calculating z-score and p-values. Network visualization was made using Gephi.

#### 3 Molecular pathways and gene ontology homogeneity (PH and GH) of cardiomyopathy-centric diseasome

In accordance with a previous study ^29^, we calculated each disease’s PH and GH homogeneity in the cardiomyopathy-centric diseasome. For PH, we used the MSigDB dataset ^58^. Each disease’s pathways homogeneity (PH) was measured as the maximum fraction of genes in the same disease with the same pathways. It is defined as:

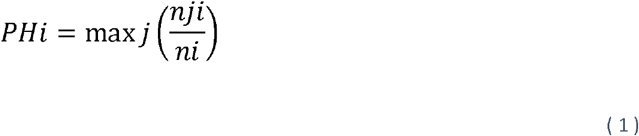

wherein *n*_*i*_ denotes the number of genes in the disorder *i* that are associated with any pathway, and *n*_*ij*_ the number of genes with a specific pathway *j*. We also created the random control PH, where the same number of genes were picked randomly from the MSigDB dataset, and pathway homogeneity was measured for them. We iterated this 10000 times and used all diseases *k* with perfect pathway homogeneity (where PH =1) for the statistical significance. The normal distribution test statistics was performed to calculate z-score:

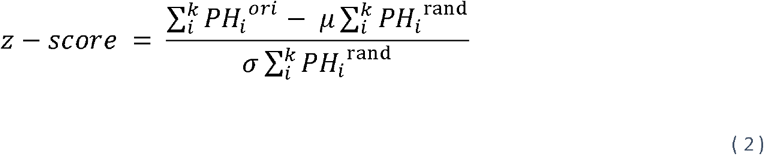

Similarly, gene ontology homogeneity analysis was applied to diseasome using the above approach. GO for disease genes were obtained from the UniProt database. This analysis mapped 4923 out of 4995 disease genes through UniProt primary gene retrieval. GH was calculated separately for biological process (BP), molecular function (MF), and cellular component (CC).

#### 4 Candidate gene prediction and filtering

The DIAMOnD algorithm was used on previous human interactome data to predict new candidate genes. The DIAMOnD algorithm uses hypergeometric distribution for new genes prediction ^32^. It is defined by Ghiassian *et. al* as:

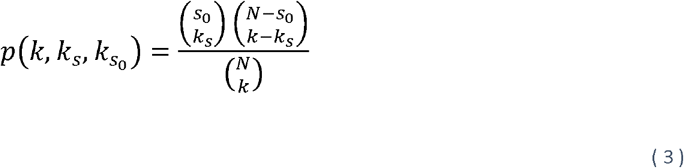

where *N* is network of proteins containing a small number (*s*_*0*_) of seed proteins associated with a particular disease and 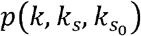 is probability that a protein with a total of *k* links has exactly *k*_*s*_ links to seed proteins. To assess whether a particular protein has more connections to seed proteins than expected under null hypothesis, cumulative probability *p*−*value* (*k,k*_*s*_) is calculated as:

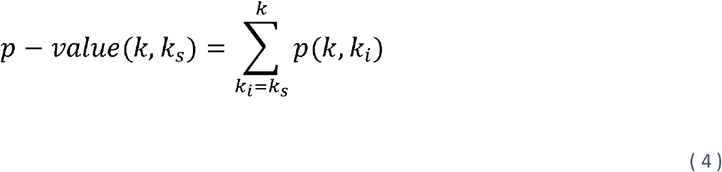

The DIAMOnD algorithm iteratively ranks all the proteins in the network. Therefore, in order to set a limit on the number of potential DIAMOnD genes, a strategy is required to define the boundary for accurate predictions. We used the MSIgDB pathways for independent biologically relevant associations ^58^. We retrieved the biological pathways significantly enriched with seed genes (Fisher’s exact test and FDR corrected using BH). Next, we assessed if these pathways were also enriched in a statistically significant manner with the DIAMOnD genes (p-value <0.05).

Next, mouse genes affecting the heart were used as screening for candidate genes. For this, the mouse cardiovascular abnormality phenotype dataset was downloaded from the International Mouse Phenotypic Consortium (IMPC), and non-heart specific phenotypes genes were dropped from further analysis ^59^. To compare with candidate genes, mouse genes were first mapped to human orthologs from the HGNC dataset ^51^. Overlapping genes between the candidates and the mouse orthologs were considered genuine potential candidates.

#### 5 Tissue specific expression of candidate genes

Tissue-specific RNA expression data was downloaded from Human Protein Atlas (https://www.proteinatlas.org/) ^39^. From this, only the GTEx dataset was used to ascertain candidate gene expression. We considered protein-transcript per million (pTPM) value greater than two as a cut-off for the presence of a gene in the tissue of interest. We also looked at the expression of candidates in the disease tissues. For this disease-specific RNA-seq datasets were downloaded from the ENA browser (https://www.ebi.ac.uk/ena/browser/home). FASTQ format RNA-seq data of study Project IDs Alzheimer’s (PRJEB28518), DCM (PRJNA494688), HCM (PRJNA352990), non-alcoholic fatty liver disease (PRJNA558102), diabetes mellitus (PRJNA358470) and rheumatoid arthritis (SRP009315) were downloaded. The raw sequences were first subjected to quality check using FastQC tool kit version 0.11.8. The pre-processed reads were aligned against human reference transcripts version 38 Ensembl release 96 using fast aligner Salmon version 2.7.3a ^60^. Downstream analysis was carried out in the R 3.4 package.

#### 6. Visualization of network and transcriptomic data

Gephi tool was used for network visualization. The circos plot was constructed using Circos software. The Heatmap and Violin plot were generated using python packages matplotlib and seaborn.

### Quantification and statistical analysis

The statistics used in the study is described in the individual method sections above.

## Acknowledgements

We would like to thank P. Murugavel, Adwait Joshi, Nitish Malhotra and Neha Kalmankar for technical help and discussions. PKC would like to thank NCBS-TIFR (Department of Atomic Energy, India) for the fellowship. The authors would like to thank NCBS (TIFR) for infrastructural facilities.

## Author Contributions statement

RS and PKC conceptualized the study. PKC collected data, performed data analysis and visualization. PKC wrote the manuscript and RS wrote the final draft of manuscript.

## Data and code availability

This paper presents an analysis of existing, publicly available data. All codes are available at the following link http://caps.ncbs.res.in/download/cardiomyo_diseasome.

Any additional information required to re-analyse the data can be acquired from the authors.

## Supplementary Information

Supplementary Materials and Figures. Further details regarding disease-genes, diseasome properties, modifier genes in each cardiomyopathy.

**Figure S1**: **Cardiomyopathy diseasome connectivity distribution, related to Figure 2**.

This figure shows connectivity distribution of cardiomyopathy diseasome. Majority of diseases are connected to a few other diseases only. DCM and HCM show high connectivity with k-value of 96 and 63 respectively.

**Figure S2: Boundary estimation of the predicted DIAMOnD genes, related to figure 4**.

Graph illustration of the biological validation of the predicted DIAMOnD genes. (A,B,C,D) Panels correspond to true molecular pathways hits and corresponding p-values of DIAMOnD genes of idiopathic cardiomyopathy (A), hypertrophic obstructive cardiomyopathy (B), restrictive cardiomyopathy (C), and histocoid cardiomyopathy (D), respectively. These pathways were firstly enriched (adjusted p-value = 0.05) using seed genes of individual cardiomyopathy. In the pathways hits plot, red lines depict seed genes and green lines refer to the DIAMOnD genes. In the p-value plot, red line highlights p-value of seed genes and green line marks the p-value = 0.05. E. Venn diagrams showing overlap between predicted DIAMOnD genes and genes showing abnormal heart phenotype in mouse for the above diseases.

**Figure S3: Interaction coverage of the human ribosomal complex proteins in reference datasets, related to Figure 4**.

Network visualization of the ribosomal complex interactions in HuRI (A), Bioplex3 (B), our human interactome (C), and STRING (D) datasets. The isolated proteins in each dataset are labelled to distinguish them in the network.

**Table S1: Disease-gene genetic data, related to Figure 1-2**.

**A**. List of the all diseases considered in this study. **B**. Various cardiomyopathies and genes implicated in them. **C**. A bipartite network data for the diseases with at least one cardiomyopathy gene. **D**. The disease projection data of the bipartite network.

**Table S2: Network properties of the cardiomyopathy diseasome, related to Figure 2**.

List of the Degree Centrality (DC), Betweenness Centrality (BC), Closeness Centrality (CC) and Clustering for each disease in the cardiomyopathy diseasome. These network statistics show the connectivity of the diseasome.

**Table S3: List of modifier genes in HCM**,**DCM and ACM, related to Figure 4**.

A List of the filtered DIAMOnD genes in the hypertrophic cardiomyopathy, dilated cardiomyopathy and arrhythmogenic cardiomyopathy.

**Table S4: Modifier genes in disease-disease association and their HPA data, related to Table 1 and Figure 5**.

**A.** List of new cardiomyopathy and other diseases association due to the modifier genes.

**B.** Major modifier genes in terms of frequency in the disease-disease association. **C**. The RNA expression (pTPM) of the modifier genes in the HPA dataset.

