## Supplemental Information for "Integrative network analysis interweaves the missing links in cardiomyopathy diseasome"

**Supplementary Figures**

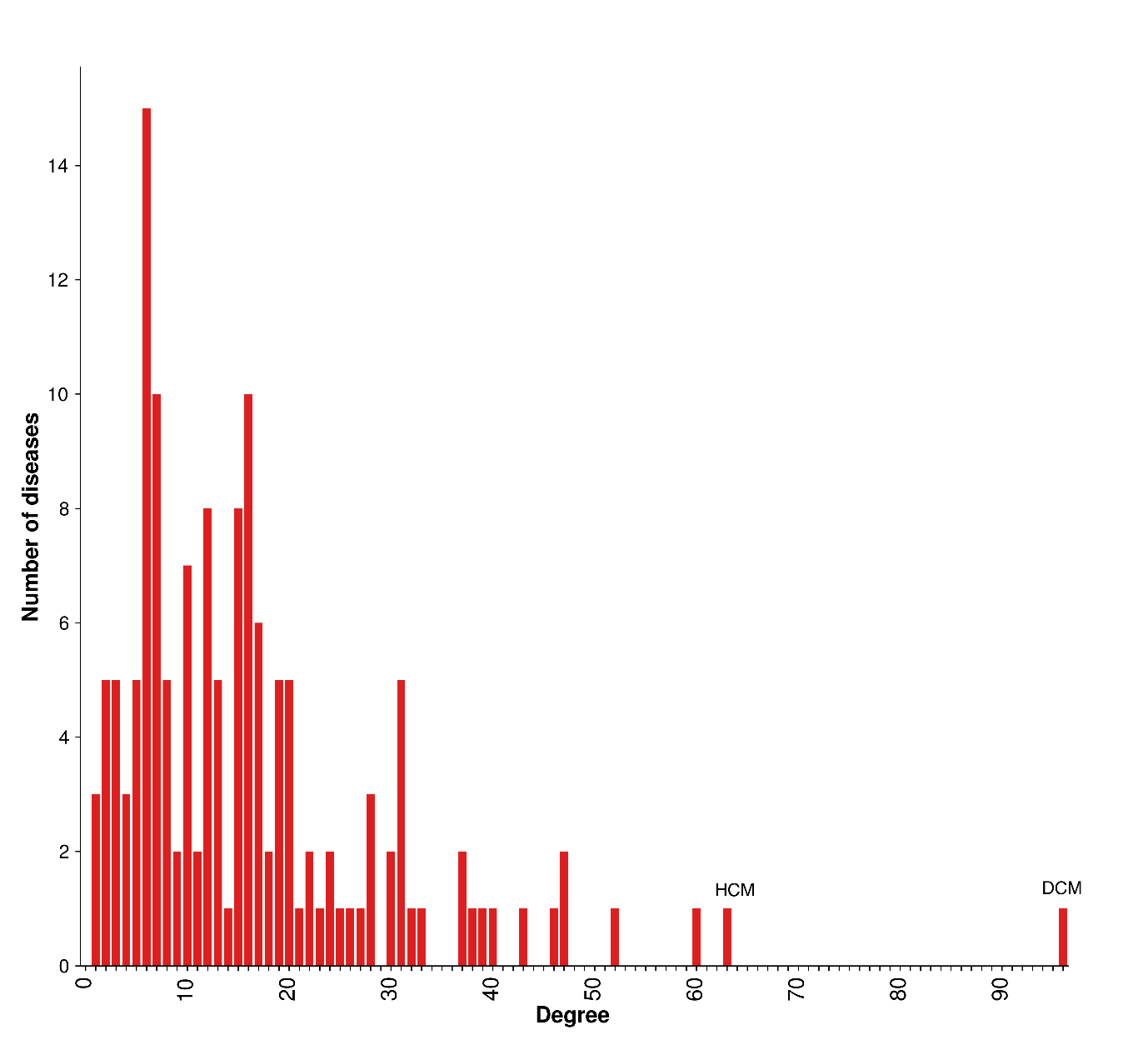

**Figure S1: Cardiomyopathy diseasome connectivity distribution, related to Figure 2.**

This figure shows connectivity distribution of cardiomyopathy diseasome. Majority of diseases are connected to a few other diseases only. DCM and HCM show high connectivity with k-value of 96 and 63 respectively.

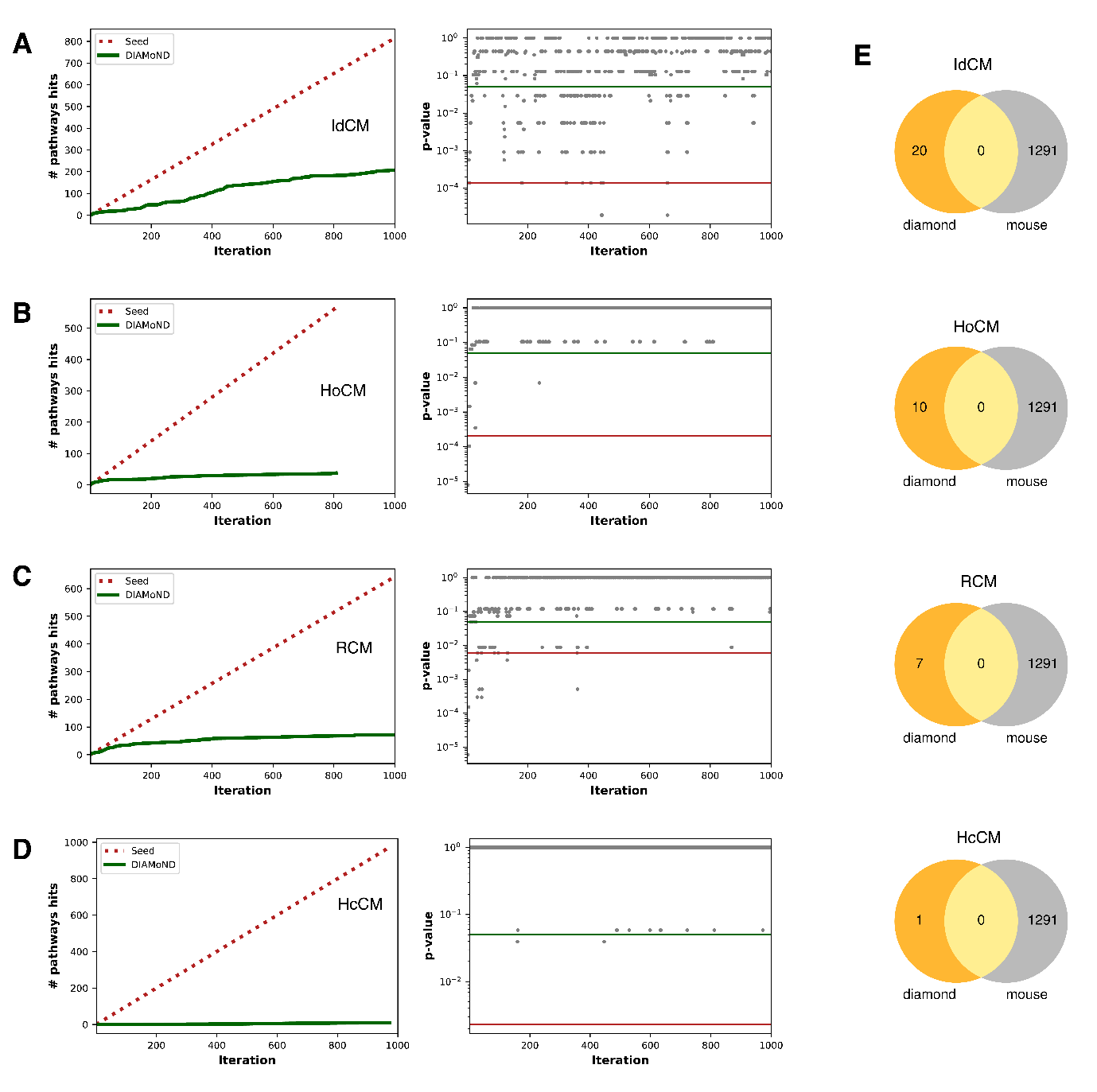

**Figure S2: Boundary estimation of the predicted** **DIAMOnD genes, related to figure 4.**

Graph illustration of the biological validation of the predicted DIAMOnD genes. (**A,B,C,D**) Panels correspond to true molecular pathways hits and corresponding p-values of DIAMOnD genes of idiopathic cardiomyopathy (**A**), hypertrophic obstructive cardiomyopathy (**B**), restrictive cardiomyopathy (**C**), and histocoid cardiomyopathy (**D**), respectively. These pathways were firstly enriched (adjusted p-value = 0.05) using seed genes of individual cardiomyopathy. In the pathways hits plot, red lines depict seed genes and green lines refer to the DIAMOnD genes. In the p-value plot, red line highlights p-value of seed genes and green line marks the p-value = 0.05. **E.** Venn diagrams showing overlap between predicted DIAMOnD genes and genes showing abnormal heart phenotype in mouse for the above diseases.

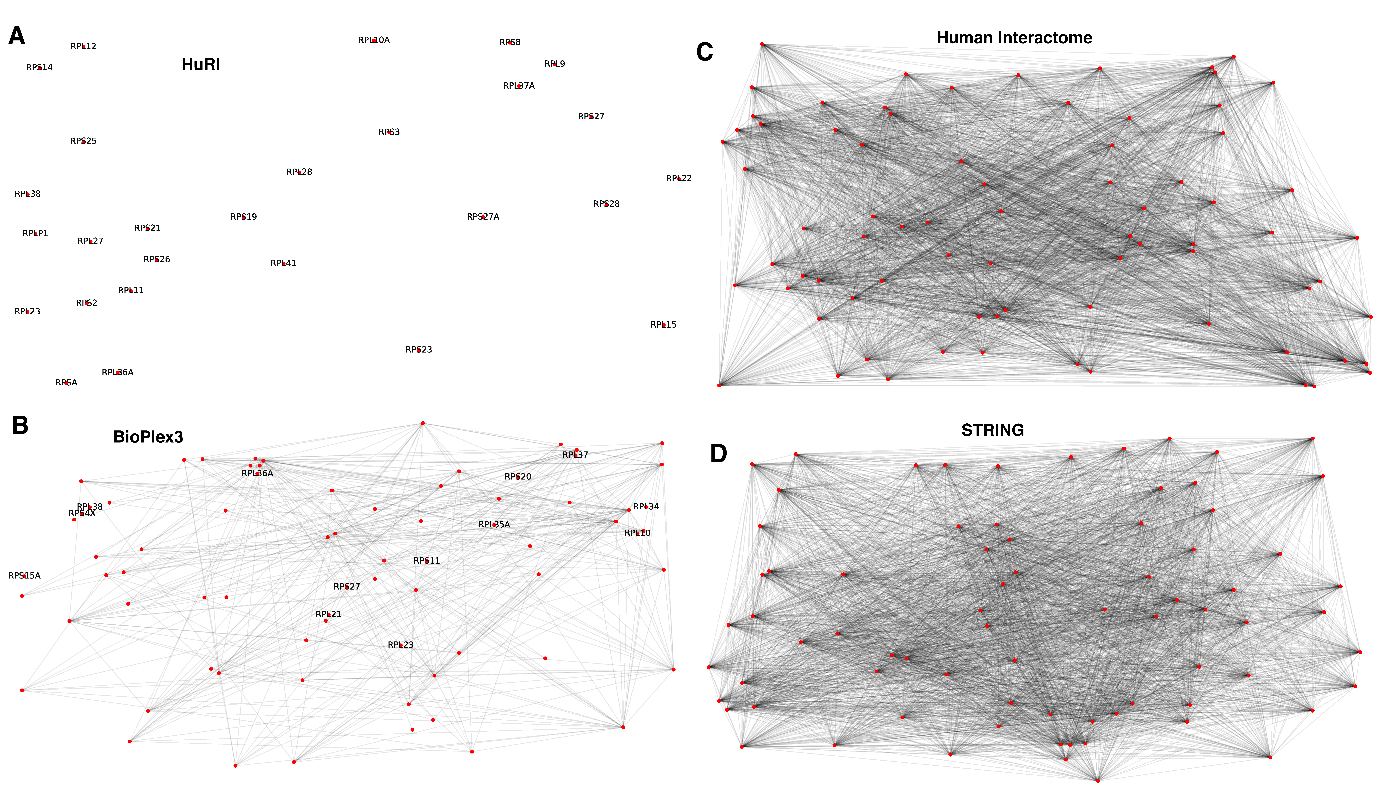

**Figure S3: Interaction coverage of the human ribosomal complex proteins in reference datasets, related to Figure 4.**

Network visualization of the ribosomal complex interactions in HuRI (**A**), Bioplex3 (**B**), our human interactome (**C**), and STRING (**D**) datasets. The isolated proteins in each dataset are labelled to distinguish them in the network.
