## Supplementary figures and images for "Integrative network analysis interweaves the missing links in cardiomyopathy diseasome"

### Figure_S1_Disease_degree_low.tif

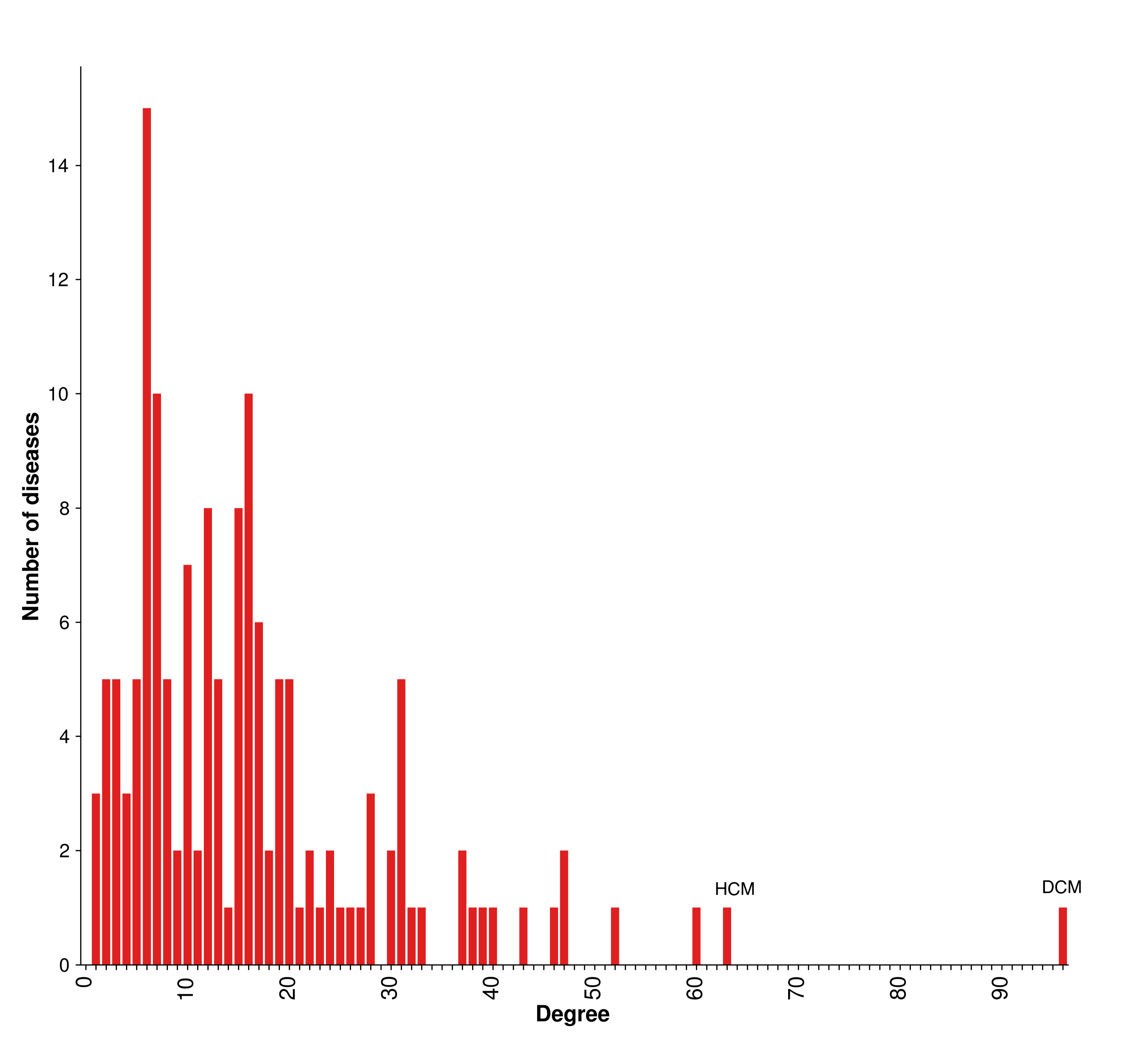

### Figure_S2_Modifier_low.tif

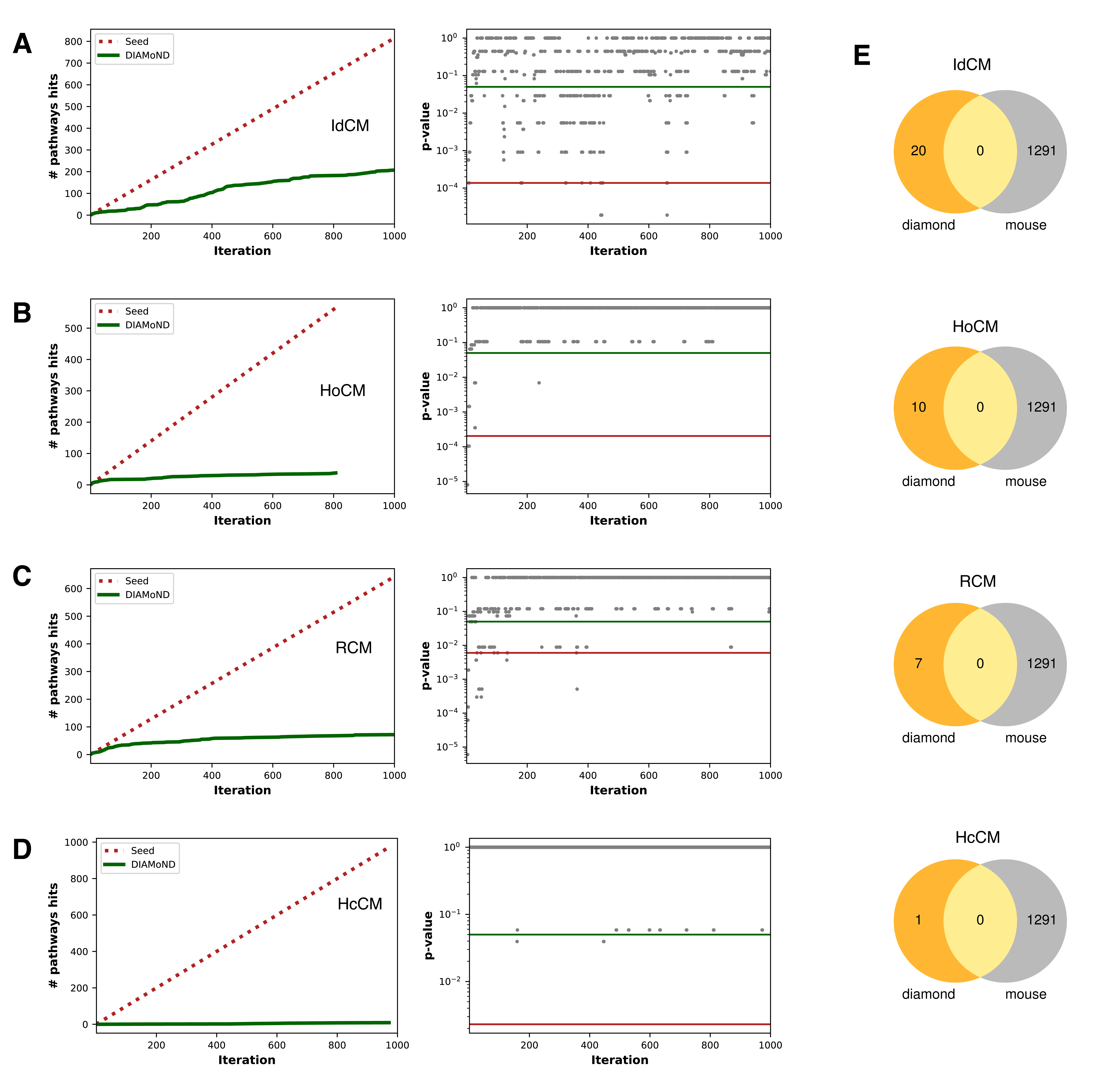

### Figure_S3_Ribosome_interactions_low.tif

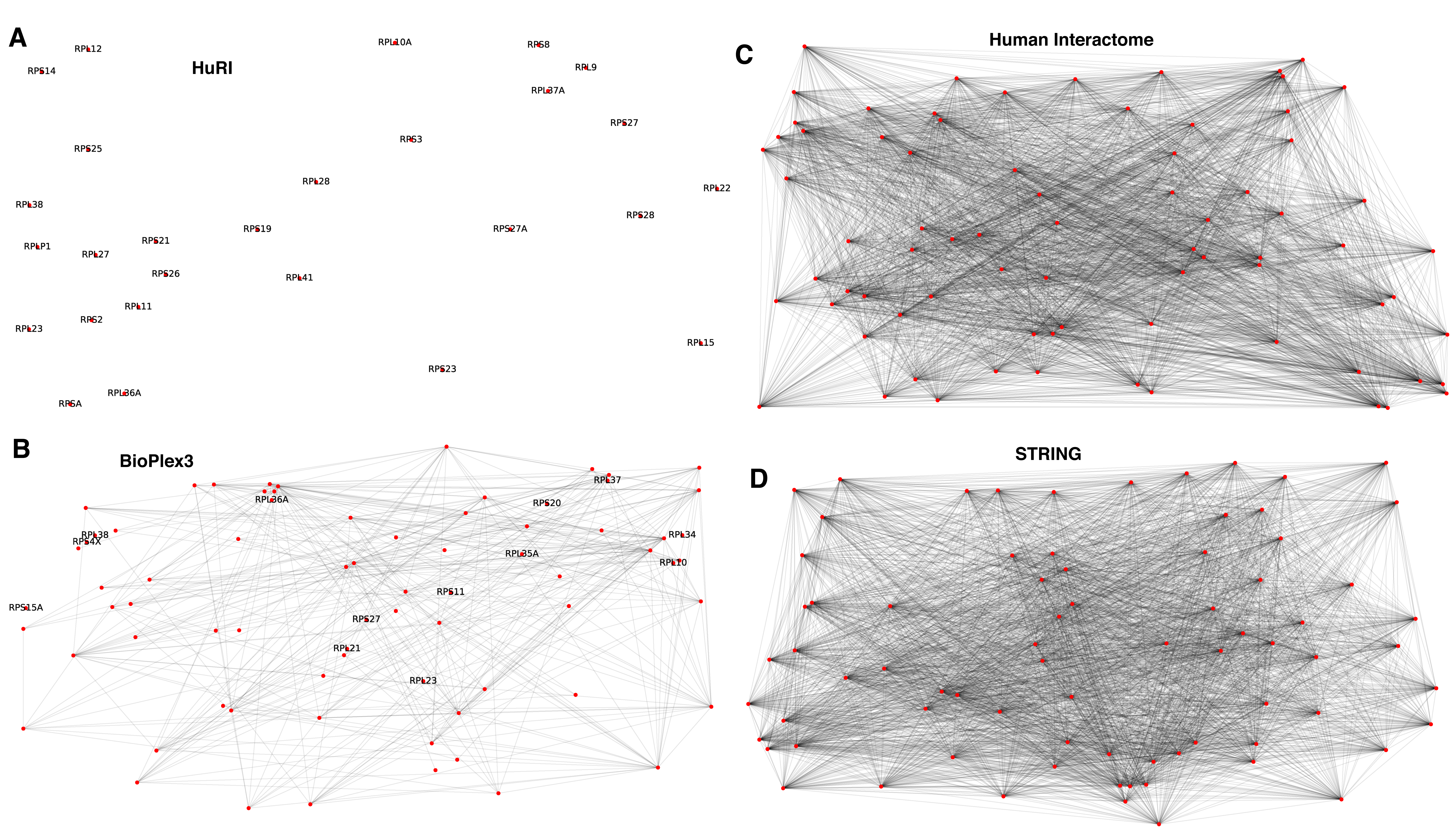
